# SynCom101: A web-based platform for the standardized design of functionally tailored synthetic microbial communities

**DOI:** 10.64898/2026.04.23.720341

**Authors:** Jiayi Jing, Siemen Rockx, Aster Liu, Chrats Melkonian, Jos M. Raaijmakers, Paolina Garbeva, Marnix H. Medema

## Abstract

**Background:** Synthetic microbial communities (SynComs) are essential tools for dissecting the causal mechanisms in host-microbiota interactions. To date, however, SynCom design suffers from a lack of standardization, typically oscillating between arbitrary strain selection and computational pipelines that misalign with experimental design. As microbiome research transitions toward functionally defined community systems with reproducible experimental outcomes, there is a strong need for a user-friendly platform that integrates multi-dimensional genomic and/or biological data into a standardized and tailored SynComs design.

**Results:** Here, we present SynCom101, a web-based platform that democratizes the design of reproducible, hypothesis-driven SynComs. SynCom101 accommodates diverse input formats including genomic annotations and laboratory-obtained phenotypic traits, allowing users to customize their design criteria with high flexibility. The platform utilizes a parsimony algorithm to ensure computational scalability for large datasets, complemented by an optional correlation-aware mode to account for microbial compatibility and co-occurrence patterns when ecological interactions among strains are available. A core innovation of SynCom101 is its suite of trait-weighting modules, which empowers researchers to strategically guide the selection algorithm toward maximal functional trait coverage, the emulation of natural community architectures, or the enrichment of positively correlated microbial assemblages to enhance community stability. We showcase the functionalities of the platform by *in silico* design of communities from different datasets, demonstrating its capacity to generate concise, functionally prioritized SynComs aligned with targeted design objectives.

**Conclusion:** By providing a transparent, parameter-documented workflow, SynCom101 ensures that community design is no longer a “black box” but a reproducible scientific record. This platform establishes a necessary standard for *in silico* community assembly, facilitating the transition from descriptive microbiome studies toward high-throughput, predictive functional screening and cross-study comparability.

**Availability:** SynCom101 can be accessed via the web interface (https://syncom101.bioinformatics.nl/). The datasets used for case studies are available on Zenodo (https://doi.org/10.5281/zenodo.18310451). The source code is available at Git (https://git.wur.nl/jiayi.jing/syncom101).

## Background

Microbial communities are fundamental regulators of host health [1–3], biogeochemical cycles [4,5], and environmental stability [6,7] across diverse ecosystems, from the human gut [8,9] to soil [10,11]. Understanding the complex, multi-species interactions within these communities is essential for advancing fields such as drug discovery [12], sustainable agriculture [13,14], and biotechnology [7,15,16]. While modern sequencing technologies have provided an expansive census of microbial diversity, translating these correlative observations into mechanistic insights remains highly challenging. This mechanistic gap is largely sustained by the inherent complexity and functional redundancy of natural microbiota, which obscure the specific contributions of individual taxa to phenotypes.

Consequently, synthetic microbial communities (SynComs) have emerged as simplified yet powerful model systems. By allowing for the precise control of community composition, SynComs provide a tractable experimental framework for testing hypotheses and uncovering mechanistic relationships governing microbial functions and interspecies and cross-kingdom interactions. Thus, the effective construction and deployment of SynComs is recognized as a decisive step from descriptive studies to predictive microbiome engineering.

Despite their utility, the potential of SynComs is constrained by a lack of methodological standardization and the inability of existing tools to support the diverse design strategies required in modern microbiome research [17]. A robust design platform must be sufficiently flexible to address a broad spectrum of hypotheses from maximizing total functional redundancy to emulating the resilience of natural communities [18], to targeting specific metabolic pathways or generating systematic community variants (e.g., N+1/N-1 community member) to probe causal interactions [19]. This need for precision is further amplified by the emergence of high-throughput phenotyping technologies, such as the kChip platform [20] and other high-throughput phenotyping facilities [21,22]. However, even these large-scale studies still predominantly rely on stochastic or random assemblies [20,23,24]. While random selection provides the “noisy” data necessary for machine learning (ML) to identify broad patterns, it is an inherently inefficient strategy for exploring the vast, non-linear functional landscapes of microbial consortia.

The future of microbiome engineering necessitates a paradigm shift from unbiased sampling of community space to the targeted design of SynComs with predefined functions. Such rationally designed communities would significantly enhance the power of ML models to pinpoint the specific traits and complementary or synergistic interactions driving host phenotypes. Furthermore, without a standardized platform to document the logic behind these designs, the assembly process remains a “black box” hindering the establishment of community standards and the comparability of results across laboratories [25].

To address this critical gap, we developed **SynCom101**, a comprehensive, web-based platform designed to standardize and accelerate the design of functionally tailored SynComs. SynCom101 provides the computational infrastructure to transition from *ad hoc* selection to the systematic generation of targeted communities. Featuring a no-code interface, the platform accommodates diverse inputs, including genomic annotations (such as KEGG annotations and biosynthetic gene clusters [BGCs]) [26–29] and experimental phenotypic traits (such as carbohydrate utilization profiles) [30,31]. This allows researchers to customize and record their selection criteria with unprecedented precision. To illustrate its functionality, we applied SynCom101 to three distinct case studies with contrasting functional prioritization strategies, demonstrating that the resulting SynComs effectively represent and maximize the targeted biological functions. Ultimately, SynCom101 serves as a foundational resource for reproducible design of functional synthetic communities, bridging the gap between high-throughput data generation and mechanistic discovery.

## Results

### Conceptual workflow of SynCom101

To standardize the transition from genomic or phenotypic bacterial datasets to functional community design, the SynCom101 workflow is structured into six integrated steps designed to maximize functional representation while ensuring experimental tractability (Fig.1A). It commences with a versatile traits-parser (Step 1) that standardizes diverse inputs into a unified, high-efficiency pickle data format. This integration ensures that the tool can accommodate both bioinformatic predictions and laboratory-obtained phenotypic data. The core of the platform is a trait-weighting system (Step 2) with customized parameter setting (Step 3) and a parsimony-based selection algorithm (Step 4). The weighting module empowers users to prioritize specific biological niches through rarity weighting or manual functional group-based adjustments, providing the flexibility to accommodate varied ecological design principles. The selection engine then identifies the best set(s) of strains required to optimize the coverage of these prioritized traits within a user-defined community size (Step 4).

**Fig. 1.**
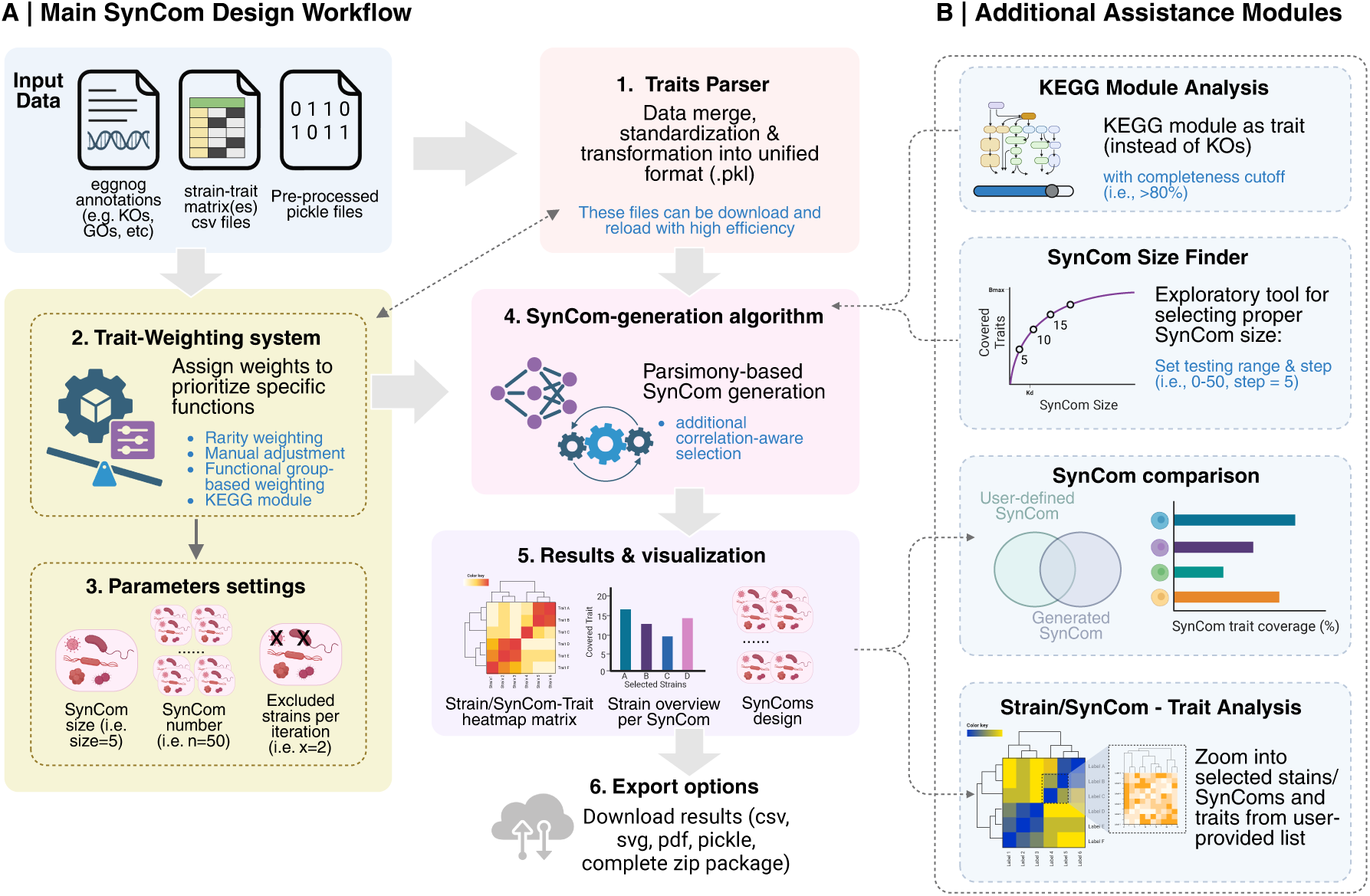
SynCom101 workflow and architecture. **(A)** Main SynCom design workflow showing six sequential steps from input data to export. **(B)** Additional assistance modules providing specialized analysis capabilities. Illustrates modular design enabling both basic and advanced user workflow.

To further refine community stability, an optional correlation-aware mode integrates strain compatibility or co-occurrence patterns, allowing for the selection of synergistic microbial assemblages (Step 4). The platform’s output includes comprehensive visualization suites that provide a landscape-level overview of the generated SynComs (Step 5). Crucially, for the sake of reproducibility, the Export module (Step 6) allows for the download of complete data packages, including high-resolution vector graphics and the precise parameter “blueprints” used for the generation, ensuring cross-laboratory repeatability and comparability.

To assist and refine the design of SynComs, the platform includes several additional modules (Fig.1B) for iterative optimization. The *SynCom Size Finder page* allows users to identify the optimal community size by simulating functional coverage across a range of sizes. The *Trait Analysis page* further allow users to compare different batch of SynComs with different design strategies and different sets of functional traits. In essence, by distilling complex microbial candidate pools into tailored consortia, SynCom101 provides a rigorous framework bridging researcher moving from *in silico* prediction to *in vitro/vivo* validation.

### Algorithmic benchmarking and computational performance

To ensure scalability for increasingly large microbial collections, we first evaluated the Traits Parser across four diverse datasets derived from eggNOG-mapper annotations. We observed that processing throughput is primarily dictated by trait density. While high-density annotations (i.e. KEGG Orthology (KOs) and Gene Ontology (GO) terms) achieved a throughput of 3–5 genomes per second, more specialized datasets like Carbohydrate-Active Enzymes (CAZy) reached 30–40 genomes per second (Fig. 2A). Across all annotation types, processing time scaled linearly with dataset size, confirming the platform’s capacity to handle thousands of genomes within minutes.

**Fig. 2.**
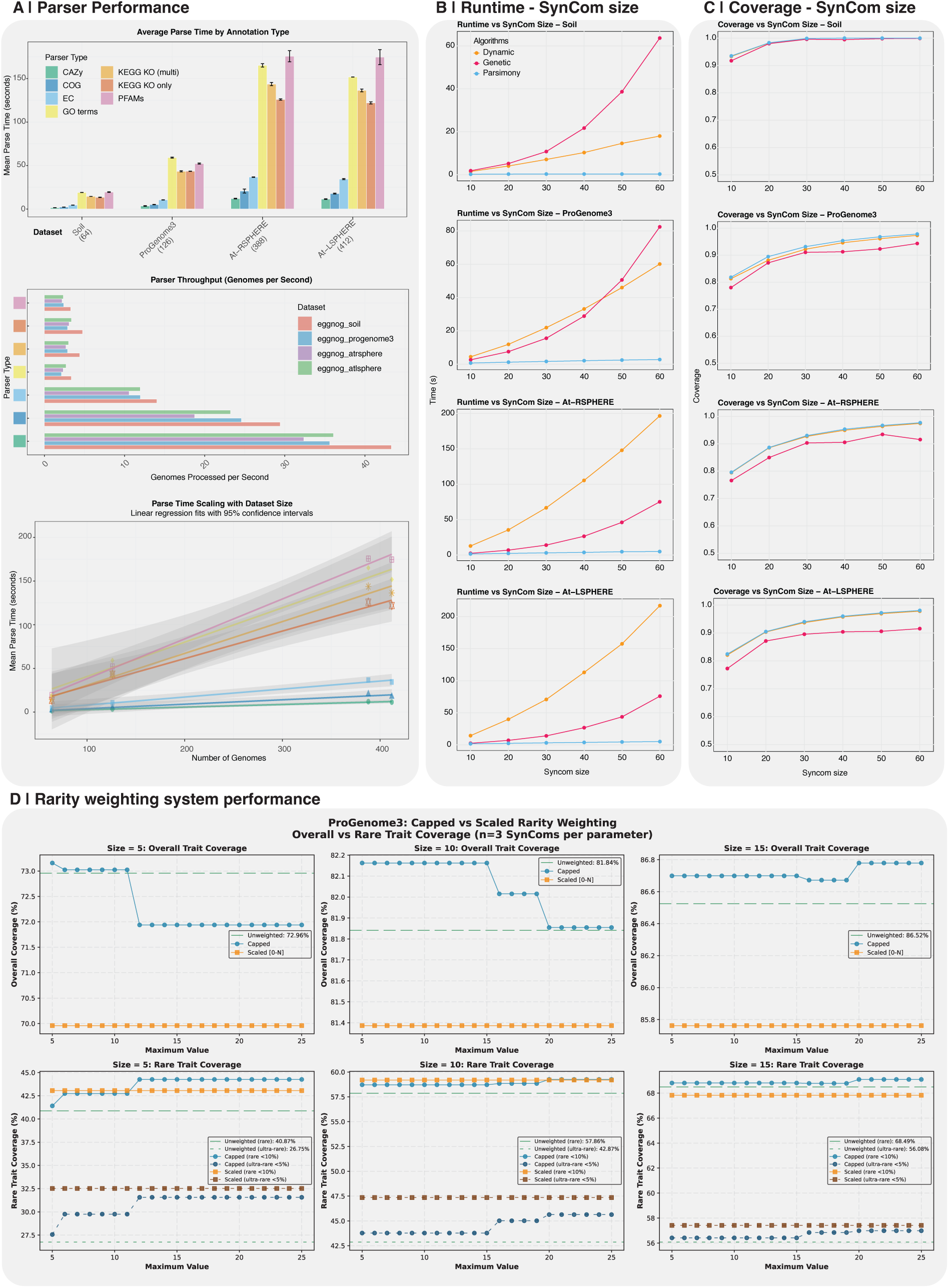
Performance benchmarks for different algorithms and SynCom design parameters. **(A)** Parser performance across annotation types and dataset sizes. **(B)** Algorithm performance comparison runtime vs SynCom size. **(C)** Algorithm performance comparison traits coverage vs SynCom size. **(D)** Rarity weighting system performance demonstrating trade-offs between overall and rare trait coverage. Traits present in less than 10% of provided strains were considered as rare traits and less than 5% were considered as ultra-rare traits.

Notably, parsing the complete annotations of 4,618 genomes (isolates genomes from crop root bacterial genome collection [CRBC] dataset) [32] from eggNOG-mapper annotations requiring approximately two hours. However, reloading these data via the standardized pickle (*.pkl*) format reduces this to less than two seconds. We therefore propose to download these exportable data through *Export* page to eliminate redundant computation and facilitate rapid, iterative design cycles.

We next benchmarked three selection algorithms (see details in Methods) to identify the optimal engine for generating SynCom with maximized functional trait coverage (Fig. 2B, C). The genetic algorithm demonstrated both the highest runtime instability and the lowest overall functional coverage across all tested datasets (Fig. 2B, C). While a dynamic programming approach yielded functional coverage nearly identical to the parsimony-based algorithm, its computational runtime increased exponentially with larger SynCom sizes (Fig. 2B). In contrast, the parsimony-based engine demonstrated superior stability and execution speed while maintaining high functional breadth, making it the ideal core for a responsive web-based interface.

Finally, we investigated the impact of functional rarity weighting using the ProGenome3 dataset, a curated collection of non-redundant, plant-associated genomes (Fig. 2D). This high-quality dataset allowed for a precise assessment of trait-capture dynamics without the noise of genomic redundancy. Our analysis revealed a critical trade-off: while “scaled” weighting (which normalizes inverse-frequency weights to a 0-1 range) maximizes the capture of ultra-rare traits (frequency <5%), it can lead to a significant reduction in overall functional diversity (Fig. 2D). Conversely, the “capped” weighting option (which imposes a mathematical ceiling on the maximum weight a rare trait can achieve) prevents extreme bias from rare traits, providing a tunable balance that enhances the retention of rare functions without compromising the general functional integrity of the consortium. This feature allows researchers to strategically calibrate SynComs to resolve specific “functional blind spots” within their candidate pools.

### Multi-scenario applications of the SynCom101 platform

The SynCom101 framework is engineered to accommodate a diverse array of ecological hypotheses, bridging the gap between large-scale microbial collections and mechanistic analysis (Fig. 3). While the Functional Specialist scenario focuses on maximizing targeted traits, the platform’s modularity allows for more sophisticated experimental designs.

**Fig. 3.**
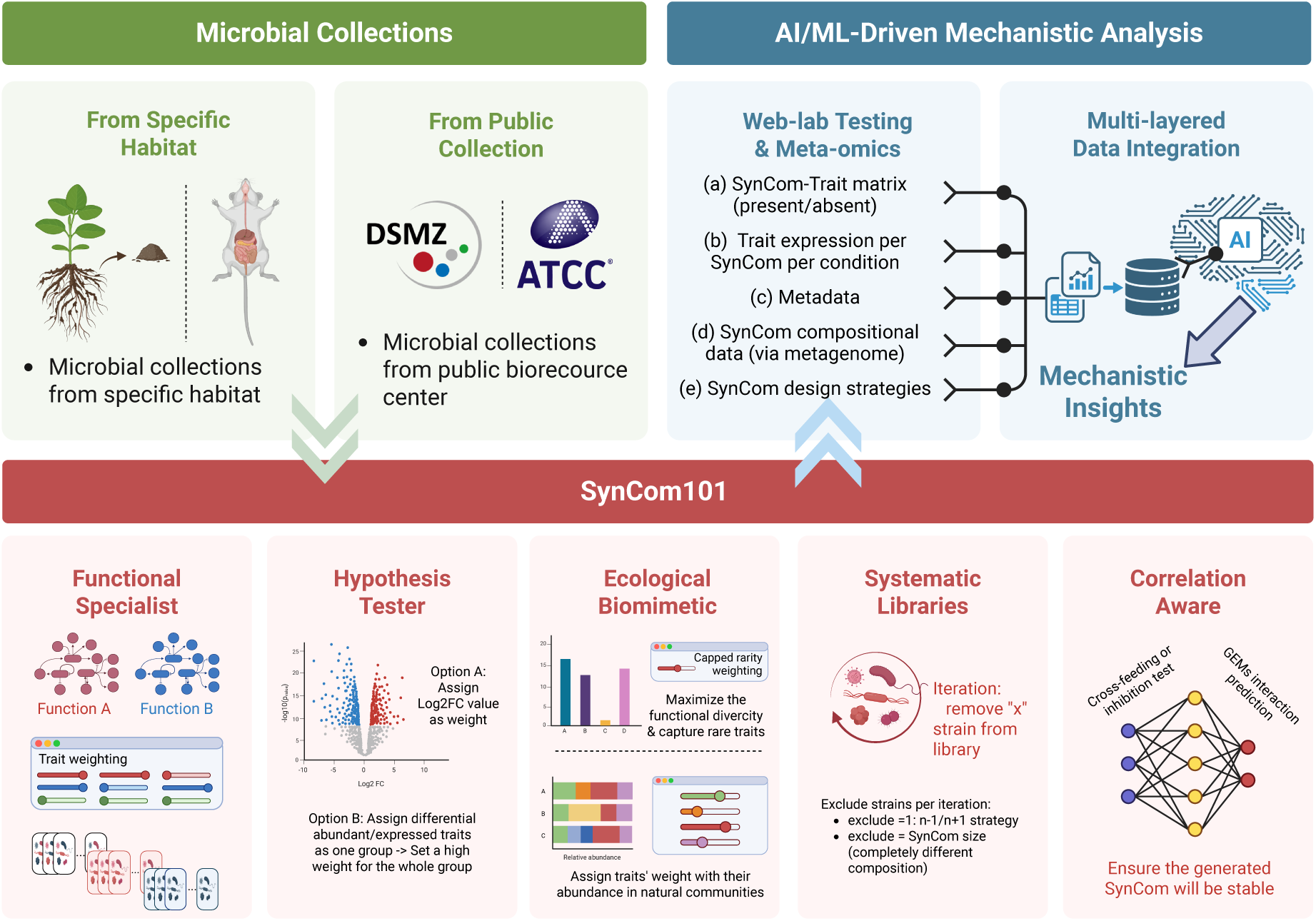
SynCom101 design framework and exemplary usage scenarios. We illustrated a conceptual workflow from microbial collection to mechanistic insights and highlighted five recommended usage scenarios for SynCom101: (1) functional specialization via targeted weighting, (2) hypothesis-driven design using experimental data (e.g., log2FC: log2FoldChange), (3) ecological biomimicry based on natural abundance patterns, (4) systematic library generation through iterative compositional shifts, and (5) stability-focused assembly via correlation-aware selection. These digital workflows facilitate downstream laboratory validation and mechanistic discovery.

In the Hypothesis Tester scenario, researchers can integrate multi-omics data (i.e. transcriptomic or proteomic profiles) by assigning selection weights based on differential expression metrics (e.g., Log2FoldChange). This strategy enables the design of “activity-centric” SynComs enriched with genes actively expressed under specific environmental stressors, facilitating the direct validation of genotype-to-phenotype predictions. To further bridge the gap between synthetic models and natural systems, the Ecological Biomimetic scenario allows for functional weighting based on *in situ* abundance patterns. This enables the assembly of consortia that recapitulate the functional architecture of native microbiomes while maintaining the tractability of a defined SynCom.

For researchers focused on causal deconstruction, SynCom101 supports the generation of Systematic Libraries. By utilizing the iterative “exclude strain” feature, users can systematically generate nearly isogenic community variants (e.g., N+1/N-1 strategies) to identify “keystone” species or develop entirely non-overlapping SynComs to maximize the breadth of functional screening. Finally, the Correlation-Aware module addresses the challenge of community stability. By integrating user-provided interaction matrices, derived from co-occurrence networks, Genome-Scale Metabolic Models (GEMs), or cross-feeding assays, the platform allows for a tunable balance between functional potency and ecological compatibility.

Collectively, these workflows position SynCom101 as a centralized hub that transforms raw microbial census data into defined experimental units, providing the standardized “design metadata” essential for multi-layered data integration and subsequent AI-driven mechanistic insights (Fig. 3).

### Example 1: rational design of functional PGP SynComs via weight prioritization

To evaluate the precision of our weight-based prioritization framework, we utilized a comprehensive genomic dataset of 4,618 crop root-associated bacteria (CRBC datasets)[32]. We targeted three core Plant Growth Promotion (PGP) categories: nutrient utilization (58 KOs), phytohormone biosynthesis (“Growth”, 22 KOs), and stress resistance (“Adaptation”, 11 KOs). By simulating distinct ecological design goals through weight adjustment, we generated 50 parallel SynComs (size n = 5) for each strategy, alongside random assembly controls to benchmark the baseline functional potential of the collection.

The strain-level analysis of single function prioritization strategies revealed a striking functional partitioning within the CRBC library (Fig. 4A). We observed that the majority of selected strains were highly specialized, with minimal overlap between “Growth” and “Adaptation” cohorts. Remarkably, only 20 strains were consistently selected in the SynComs sets across three single-function SynCom design criteria. This scarcity of “generalist” strains underscores the inherent difficulty in finding single isolates that harbour a complete PGP repertoire and reinforces the necessity of a rational SynCom approach to achieve multi-functional synergy in the rhizosphere.

**Fig. 4.**
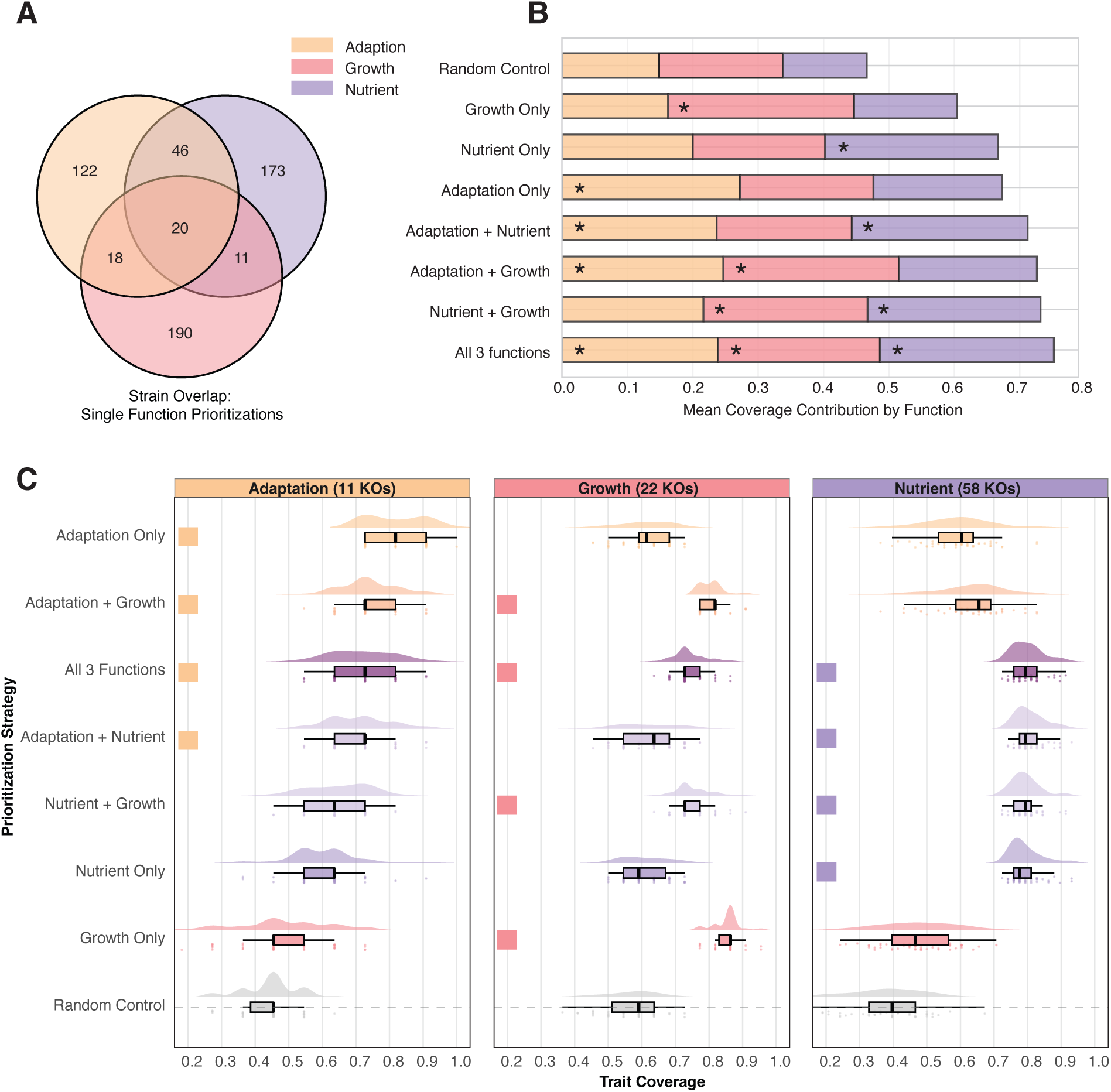
Example SynComs design with CRBC dataset with plant growth promotion traits prioritized. Each strategy involved 50 SynComs with 5 strains per SynCom. <u>Random Control</u>: random selected SynComs as control. <u>Growth only</u>: 22 plant growth related KOs were set weight = 1, all other KOs weight = 0. <u>Adaptation/Nutrient Only</u>: similar to ‘Growth only’ strategy, the related KOs were set weight = 1, all other KOs weight = 0. <u>Adaptation + Nutrient</u>: both KOs were prioritized. <u>All 3 functions</u>: all 3 PGP related functions were prioritized. The full PGP-KOs list sees supplementary material in CRBC paper [32]. **(A)** Strain overlap Venn diagram for three single-function prioritization design strategies. **(B)** Mean coverage contribution by function across prioritization strategies. The “*” highlighted the prioritized function(s). **(C)** Trait coverage distributions showing precision of functional targeting. The colored squares highlight the strategies the specific functions were prioritized for in the design.

Our weight-based algorithm effectively navigated these genomic constraints to enrich for target traits in the assembled communities. The mean coverage contribution for each prioritized function increased when assigned a higher weight, confirming the tool’s ability to steer community composition toward specific metabolic functions (Fig.4B). Detailed distribution analysis across the 50 parallel SynComs further highlighted the influence of genomic abundance on design precision (Fig.4C). While nutrient-focused SynComs reached a high and stable functional coverage (>70%), strategies targeting adaptation modules showed greater variance and lower median coverage. This discrepancy demonstrates that our prioritization framework provides an honest representation of the functional limits of the available microbial functional assignment. Critically, this entire iterative workflow is fully integrated within SynCom101. Users can seamlessly transition from the design module to the Trait Analysis sub-page, where the specific functional subsets of generated SynComs are automatically extracted and statistically visualized.

### Example 2: balancing specialized traits with community-wide functional diversity

To investigate the flexibility of our prioritization framework in complex ecological contexts, we applied SynCom101 to a dataset of 356 isolates from a disease-suppressive soil (S11) [33–35]. We focused on their Biosynthetic Gene Clusters (BGCs), clustered into Gene Cluster Families (GCFs) via BiG-SCAPE to represent the strains’ metabolic potential. Given their established role in soil-borne pathogen suppression, metallophore-related GCFs were manually curated as primary functional targets [34]. We designed five distinct assembly strategies (size n=20, optimized via the ‘Size Finder’ module) to test how varying weight ratios between metallophore traits and the broader BGC landscape influence community functional profiles.

Our results demonstrate that subtle shifts in weighting factors allow for the fine-tuning of community “multi-tasking” capabilities (Fig. 5A). When metallophore traits were prioritized with high intensity relative to other BGCs (Weight=10 vs. Weight=1), the resulting SynComs not only maximized the recruitment of metallophore BGCs but also maintained a high overall BGC richness (Fig. 5B). In contrast, an “exclusive” prioritization strategy (Metallophore Weight=1, Others=0) led to a significant “functional collapse” in non-target categories, where the gain in metallophore coverage was offset by a drastic reduction in total GCF diversity (decreasing from around 600 to 379 total traits).

**Fig. 5.**
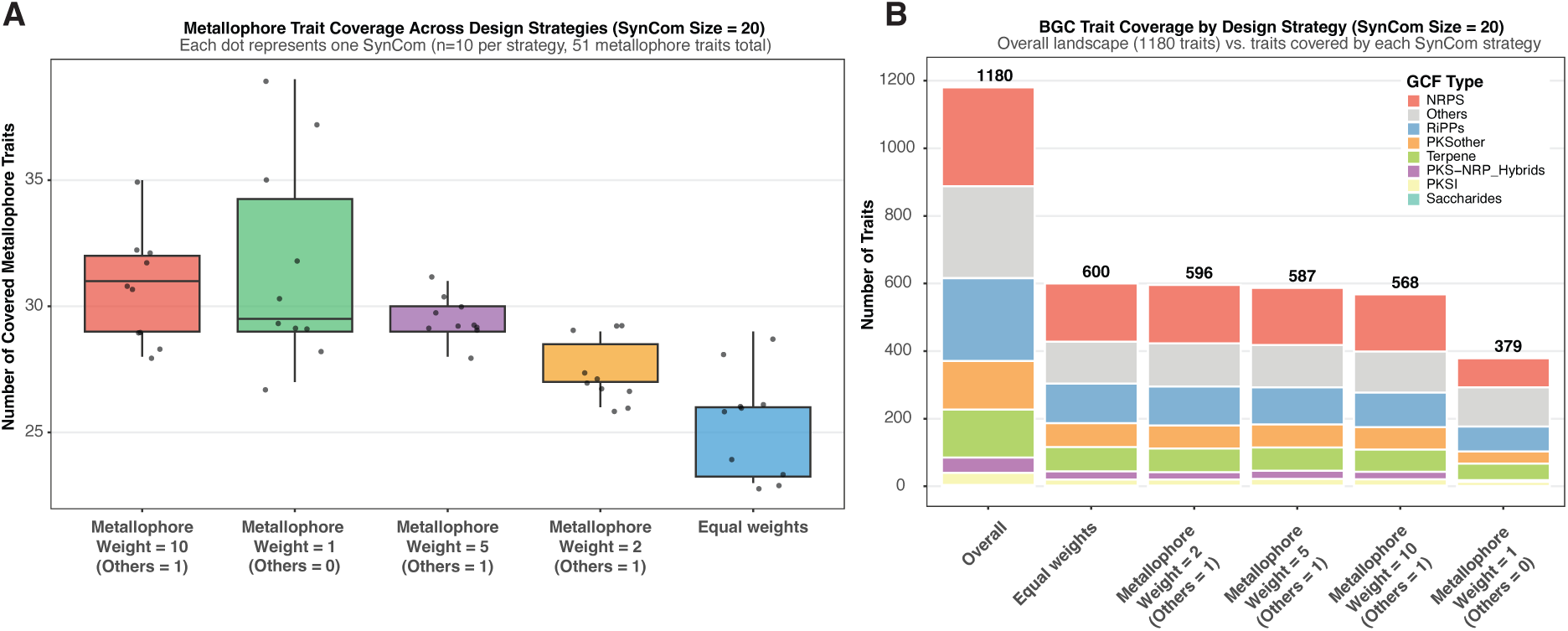
Example SynComs design with disease-suppressive soil dataset with biosynthetic gene clusters (BGCs) prioritized and metallophore BGCs with different weights. **(A)** Metallophore trait coverage across different weighting strategies (n=10 SynComs per strategy). **(B)** BGC trait coverage landscape with different BGC types.

These findings highlight the sophisticated control SynCom101 offers for navigating the trade-offs between a primarily desired functional trait and general functional potential. By enabling users to calibrate the “selective pressure” across multiple trait categories, the platform facilitates the design of SynComs that are not only potent in their targeted function but also preserve the diversity of secondary metabolic pathways. This iterative optimization process is streamlined within the platform, allowing researchers to find the balance tailored to their specific environmental application.

### Example 3: balancing cross-feeding prediction with bacLIFE-based global traits

To evaluate the correlation-aware selection mode of SynCom101, we again applied the platform to the 356-strain disease-suppressive soil collection (S11), this time annotated with bacLIFE-derived functional traits (64,904 traits in total, matrix sparsity 94.6%). To quantify metabolic complementarity, a directed cross-feeding matrix was constructed from genome-scale metabolic models (GSMMs) using gapseq [36], encoding the number of cross-feeding compounds (one strain can produce and another one can consume) between each strain pair. Five parallel design strategies were generated by systematically varying the cross-feeding weight parameter from 0 (pure function-based optimization) to 1.0 (pure correlation-based design) in four increments (0, 0.2, 0.5, 0.8, and 1.0). All other settings were held constant: 20 SynComs per condition, community size of 20 strains, five strains excluded per iteration, and automatic rarity weighting with a capped value of 10.

Increasing the cross-feeding weight produced a pronounced and monotonic trade-off between functional coverage and community cohesion (Fig. 6A). Under pure coverage optimization (weight = 0), SynComs achieved a mean weighted coverage score of 0.501 ± 0.057 with a mean cohesion score of 34.2 ± 6.4 and a mean positive cross-feeding pair fraction of 81.9%. Introducing a modest cross-feeding weight of 0.2 left coverage nearly unchanged (0.509 ± 0.049) while marginally improving cohesion (37.4 ± 6.9) and the positive pair fraction (84.1%), suggesting that a light crossfeeding bias can be applied at virtually no functional cost. As the weight was raised to 0.5, cohesion climbed substantially to 57.4 ± 5.3 and the positive pair fraction reached 94.0%, accompanied by only a moderate reduction in coverage (0.475 ± 0.037). Stronger cross-feeding weights further amplified cohesion, peaking at a mean of 78.2 ± 6.3 at weight = 1.0 but at the expense of a marked bacLIFE functional trait coverage decline (0.278 ± 0.032), representing an approximately 44% reduction relative to the unweighted baseline. Jaccard similarity analysis of strain compositions confirmed that community membership progressively diverged with increasing cross-feeding weight (Fig. 6B), indicating that the algorithm systematically recruits a distinct, metabolically compatible subset of the microbial candidates’ pool when cohesion is prioritized. The strain-level selection frequency heatmap (Fig. 6C) further revealed that certain strains became dominant under high cross-feeding weights, reflecting their central role as metabolic hubs within the pairwise interaction network.

**Fig. 6.**
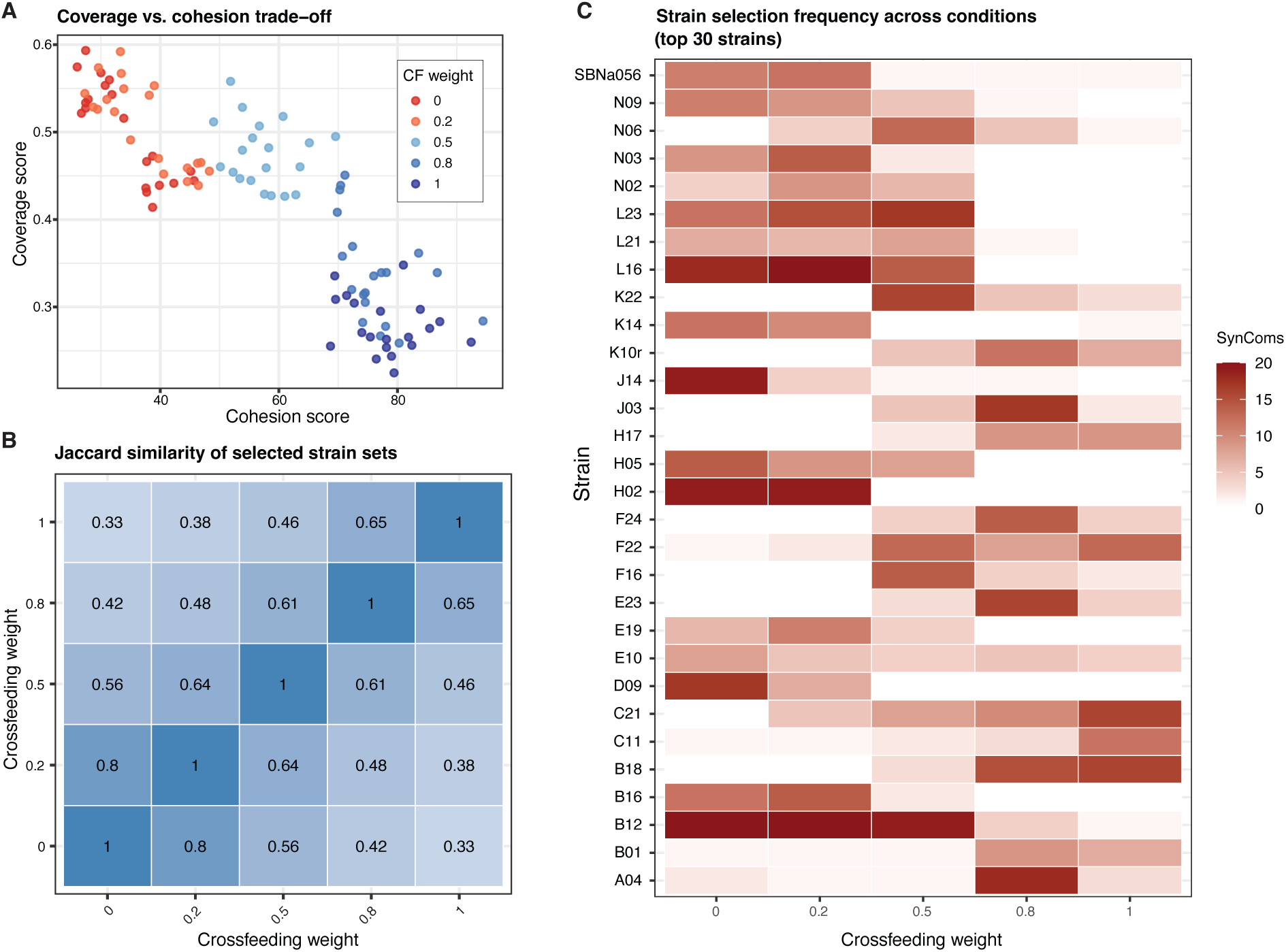
Example SynCom design with disease-suppressive soil dataset with bacLIFE clustered traits auto rarity weighting and cross-feeding prediction with GSMMs. **(A)** Trait coverage scores and cohesion scores across different weighting strategies (n=20 SynComs per strategy, SynCom size = 20, exclude per iteration = 5, auto rarity weighting capped value = 10). **(B)** Jaccard similarity of selected SynComs compositions across different cross-feeding weight setup. **(C)** Top 30 frequently selected strains across different cross-feeding weight design strategies. The heatmap color shows which specific strain were selected in how many SynComs under each cross-feeding strategy.

These results demonstrate that SynCom101’s correlation-aware mode provides precise, continuous control over the balance between functional breadth and metabolic complementarity. By adjusting a single parameter, researchers can design SynComs spanning the full spectrum from maximally diverse-functional consortia to highly cohesive, cross-feeding-optimized communities. Crucially, the intermediate weight range (0.2 - 0.5) offers a practical “optimal” zone in which substantial gains in metabolic integration are achievable with minimal sacrifice of global trait coverage. This tunability is particularly valuable when the experimental aim shifts from comprehensive functional representation toward community stability or when metabolite-exchange dynamics are hypothesized to underpin the ecological function of interest. Collectively, these findings establish SynCom101 as a flexible platform that can incorporate diverse types of inter-strain interaction data, whether derived from GSMMs, co-occurrence networks, or empirical cross-feeding assays, to assemble communities that are both functionally rich and metabolically coherent.

## Discussion

The rational design of synthetic microbial communities (SynComs) has long been hampered by the lack of a dedicated, systematic framework that can translate complex genomic potential into experimental blueprints. Historically, researchers have relied on *ad hoc* strain selection or simple taxonomic diversity to assemble communities, which is both labor-intensive and unpredictable. While recent theoretical frameworks, such as MIMIC2 [18], have begun to utilize meta-omics-guided trait prioritization to inform design, the practical implementation of these strategies still often requires bespoke bioinformatics pipelines that remain inaccessible to many experimentalists. SynCom101 fills this critical gap as a purpose-built design engine, by enabling microbiologists to translate genomic potential into experimentally testable community designs. By integrating multi-trait prioritization with a user-friendly, web-based infrastructure, the platform replaces the current reliance on bioinformatic scripts with a standardized, reproducible workflow.

Crucially, the utility of SynCom101 is further amplified by its integration of curated public bacterial collections. By hosting pre-processed genomic data from established repositories (e.g., the At-SPHERE, CRBC, metaTraits, and Progenome3, etc), the platform allows the community to repurpose existing strain libraries for diverse, novel applications. This feature transforms static collections into dynamic assets that can be iteratively re-designed and validated across different habitats, ensuring that as the global census of culturable microbes grows, our capacity to architect them into functional ecosystems scales accordingly.

The core advantage of SynCom101 lies in its operational flexibility, which the selection criteria can be customized for different research objectives and documented in a standard way. Across the three case studies, SynCom101 demonstrates its capacity to resolve key biological trade-offs, from identifying functionally specialized versus generalist strains in plant growth-promoting communities, to balancing targeted biosynthetic traits with overall metabolic diversity, and to quantitatively tuning the balance between cross-feeding-driven cohesion and global functional coverage. Unlike traditional binary selection methods, our framework utilizes a weighting system that allows users to navigate the “Specialization-Diversity” trade-off with mathematical precision. Our results from the CRBC and disease suppressive soil rhizobacteria datasets demonstrate that an exclusive focus on a single functional trait often leads to a loss of broader metabolic redundancy that is essential for community resilience. SynCom101’s ability to calibrate the “selective pressure” across multiple trait categories ensures that the resulting SynComs do not only cover a single category of targeted functions (e.g., enriched in metallophore BGCs) but are also more likely ecologically robust. This level of intentionality in design is a fundamental shift from the “trial-and-error” assembly strategies that dominate the field today.

The modularity of the platform also facilitates the exploration of complex ecological theories, such as the metabolic division of labour and niche complementarity. Through the correlation-aware and ecological biomimicry modules, researchers can move beyond simple presence/absence models to design communities that recapitulate the synergistic interactions and stability of native microbiotas. As high-throughput microfluidic technologies and robotic biofoundries continue to scale, the requirement for standardized architectural platform becomes even more acute.

Beyond algorithmics, the systematic generation of multiple community replicates with shared functional profiles offers a robust solution to the high rate of false-positive phenotypic associations in microbiome research. Traditionally, studies have relied on one or two representative SynComs to validate a functional hypothesis—a strategy that lacks the statistical power to distinguish between essential functional drivers and coincidental phenotypic observations. SynCom101 enables a more rigorous approach by facilitating the assembly of functional cohorts: groups of distinct communities designed to target the same metabolic or ecological objective. By comparing multiple, compositionally diverse SynComs optimized for Function A against a similar cohort for Function B and a set of randomly assembled controls, researchers can move toward a statistical assessment of functional importance. If a consistent phenotype is observed across a majority of communities in a functional cohort, the likelihood that the trait is driven by the targeted genes rather than individual strain idiosyncrasies is significantly higher. This ability to generate diverse, high-replicate community variants effectively mitigates the risk of anecdotal results and provides a standardized pathway for pinpointing the specific microbial traits that govern host-phenotype interactions.

Furthermore, SynCom101 introduces a level of methodological transparency that has been largely absent from the microbiome research field. Currently, the selection logic of most SynComs is documented in non-standardized prose, which hinders cross-study comparability and reproducibility. By generating exportable, parameter-documented blueprints, SynCom101 transforms community design into a standardized digital record. This standardization is essential as the field moves toward integrated meta-analyses of SynCom data, where the ability to replicate the exact selection logic across different laboratories is paramount.

In conclusion, SynCom101 democratizes the rational design of microbial ecosystems. By lowering the computational barrier for biologists while maintaining mathematical rigor, it provides the foundational infrastructure necessary to bridge the gap between the genomic census of high-throughput sequencing and the mechanistic discovery.

## Methods and Implementation

SynCom101 (https://syncom101.bioinformatics.nl/) is developed as a cloud-based platform for designing microbial synthetic communities based on genomic annotations and/or phenotypic data. It utilizes the Python programming language for backend and Streamlit (https://streamlit.io/) for frontend. The platform is organized into nine functional subpages that facilitate a comprehensive workflow from raw data parsing to results export and analysis: (1) Home page, (2) Data input, (3) Trait weighting, (4) Trait analysis, (5) Find size, (6) KEGG Analysis, (7) SynComs generation, (8) Results, and (9) Export.

### Input data format and parsers

The data ingestion layer of SynCom101 is engineered to convert diverse inputs into a standardized, internal strain-trait matrix (*syncom101_parser.write_data_matrix*). The platform supports four primary input methods, each managed by specialized backend parsers for normalization of data format:

#### Direct serialization (syncom101_parser.py)

For users with pre-existing SynCom101 projects, the platform supports the direct upload of .*pkl* or *.txt* pickle files. The *read_data_matrix* function deserializes these objects into three components: a dictionary-based functional matrix, a list of strain identifiers, and a nested dictionary of trait attributes (names, weights, and annotations).

#### Manual tabular input (syncom101_parser.py)

This module handles standard CSV or XLSX files. It allows users to define the nature of the traits using a type-mapping system (TP_MAPPING): binary presence/absence (pa) or numerical values (vl). The parser performs automated data cleaning, including the alignment of strain names across experimental and metadata tables.

#### eggNOG annotation parsers

Including syncom101_eggnog_parser.py and syncom101_optimized_multi_annotations_parser.py. This workflow ingests outputs from the eggNOG-mapper pipeline [26,37]. The platform supports the upload of individual “*.emapper.annotations*” files or bulk ZIP archives. The optimized multi-annotation parser allows for the simultaneous extraction of multiple genomic features, including KEGG Orthologs (KOs), PFAM domains, CAZy families, COG categories, Gene Ontology (GO) terms, and preferred gene names, by processing the raw annotation files in a single computational pass.

#### bacLIFE parser (syncom101_bacLIFE_parser.py)

To integrate specialized metabolism into community design, SynCom101 provides a dedicated parser for the bacLIFE workflow [29]. It ingests the “MEGAMATRIX_renamed.txt” file to extract Gene Cluster Families (GCFs). This module enables the prioritization of strains based on their secondary metabolite biosynthetic potential, mapping specific chemical signatures identified through comparative genomics into the functional design matrix.

#### Advanced KEGG module parser (syncom101_kegg_module_parser.py)

The parser applies a user-defined threshold (*minimal_score*, default 80%) to transform continuous completeness data into a binary matrix, allowing community selection based on complete metabolic pathways. This will consider complete KEGG modules instead of KOs as individual traits. It supports two technical workflows:

1. Direct Parsing: It processes outputs from *kegg-pathway-completeness* tools using regular expressions to extract KEGG Module identifiers (e.g., M00001) and their percentage completeness.
2. KO-to-Module Conversion (in KEGG analysis page): The platform can dynamically map gene-level KO data to the KEGG pathway hierarchy. It transforms continuous completeness scores into a binary or numerical functional matrix based on a user-defined threshold (*minimal_score*). This allows the selection engine to target functional redundancy or metabolic gaps at the pathway level.

### Trait weighting and prioritization

The transition from a raw strain-trait matrix to a hypothesis-driven design is governed by the assignment of weight vectors (*w_j_*) to functional traits. SynCom101 implements multiple weighting workflows that allows users to prioritize functional coverage according to specific usage scenarios.

#### Automatic rarity weighting system

To prevent the optimization engine from overlooking low-frequency traits that may be ecologically critical, the platform implements a rarity-prioritization module. The algorithm automatically calculates the frequency (*f_j_*) of each trait across the entire strain library and assigns weights using an inverse-frequency model:

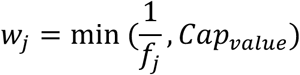

Technically, the platform allows users to define a capping value (e.g., 5). This constraint prevents ultra-rare traits from receiving disproportionately high weights that would otherwise force the selection of low-utility strains, thereby maintaining a balance between rarity preservation and total functional coverage.

#### Template-driven manual weighting

For precise control over individual traits, the platform automatically generates a customized two-column CSV template (*trait_id, weight*) based on the specific traits present in the current input data. This ensures that all unique identifiers are correctly mapped. Users download this session-specific template, edit the weight values locally, and re-upload the file. The backend performs a hash-mapping operation to synchronize the uploaded values with the internal matrix, allowing for the targeted prioritization of specific genomic or phenotypic markers.

#### Categorical functional-group weighting

This module enables strategy-level design by grouping individual traits into biological categories (e.g., “Siderophores”). The implementation follows a two-stage logic:

1. Categorical Mapping: The system generates a template where users assign each trait_id to a functional_group.
2. Dynamic UI Interactivity: Upon re-uploading the mapping, the platform dynamically renders numeric input fields in the web interface for each identified group. Users set the weight for the entire category via the UI, and the backend propagates this value to all constituent traits. This decoupling of mapping (via file) and weighting (via UI) allows for rapid iterative balancing of different functional “themes”.

#### KEGG-Specific regex weighting

For genomic datasets, a specialized interface utilizes regular expressions to filter and weight traits. By inputting keywords or metabolic identifiers (e.g., “M00001” or “transporter”), the system performs a string-match across the trait attributes and applies user-specified weights collectively to all matching indices.

### Traits-based SynComs design algorithms

SynCom101 is engineered to generate synthetic communities based on defined prioritized functions and annotations from microbial candidate’s pool provided by users. While multiple optimization paradigms including dynamic programming and genetic algorithms were benchmarked for performance, the iterative parsimony (greedy) algorithm was selected as the production engine for the web interface due to its superior computational efficiency and scalability with high-dimensional genomic datasets.

#### Parsimony engine

The core selection logic utilizes an iterative heuristic that picks the “best” strain at each step. In every iteration, the algorithm calculates a Selection Score (*S*) for all remaining strains *i* based on their weighted contribution to the traits not yet covered by the community:

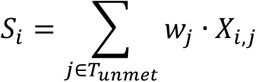

Where *X_i,j_* represents the presence (or value) of trait *j* in strain *I*, and *w_j_* is the user-defined (or rarity-adjusted) weight. The strain with the highest score is added to the SynCom, and the “unmet traits” list (*T_unmet_*) is updated before the next selection round.

#### Redundancy control and penalty mode

To allow users to design lean communities without excessive functional overlap, the platform implements an optional penalty mode. When active, the score of a candidate strain is penalized by a factor (P) for every trait it possesses that is already present in the existing community:

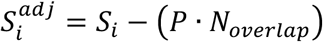

This forces the algorithm to prioritize complementary strains that fill functional gaps rather than redundant “generalist” strains.

#### Correlation-aware selection logic

To account for ecological compatibility and strain-strain interactions, SynCom101 implements a correlation-aware scoring module. This feature allows the integration of external interaction data, such as co-occurrence networks, competition assay results, or metabolic modeling predictions (e.g., from Genome-Scale Metabolic Models), directly into the iterative selection process.

1. Pairwise interaction scoring (syncom101_correlation_scoring.py): During each selection step, every candidate strain (*s*) is evaluated against the set of strains already included in the current SynCom (*C_current_*). A Correlation Bonus (*B_s_*) is calculated as the sum of weighted pairwise interactions:

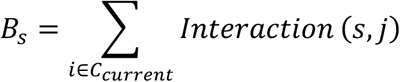 Technically, the module applies a positive weight to synergistic interactions (correlations \gt 0) and a negative penalty to antagonistic interactions (correlations \lt 0). This ensures that the algorithm avoids co-selecting strains that are likely to inhibit one another in a laboratory or natural environment.
2. Crossfeeding matrix processing (syncom101_correlation_scoring.py): When using genome-based metabolic models (GSMMs)-derived predictions, the users upload a directed crossfeeding matrix in which entry *matrix_i,j_* encodes the number of metabolites that strain *i* can produce and strain *j* can consume. To capture bidirentional metabolitc exchange potential, the raw asymmetric matrix is symmetrized by summing it with its transpose, and the resulting values are normalized to [0,1] by dividing by the global maximum of the symmetrized matrix. The diagonal is set to zero to exclude self-crossfeeding. This normalized symmetric matrix input from a computational standpoint while preserving the metabolitc interpretation of the data.
3. Multi-objective optimization function: When correlation mode is active, the final selection score for a strain (*S_total_*) is a weighted balance between its functional contribution and its ecological compatibility:
 

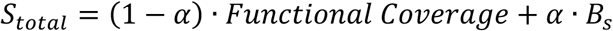 The correlation weight (α), adjustable via the web interface (0.0 – 1.0), allow users to prioritize either maximum functional breadth (low α) or maximum community stability/cohesion (high α).
4. Post-selection cohesion metrics: After SynCom generation, the platform calculates an interaction summary to validate the design. For correlation-based designs, this includes: <u>(a)</u> positive/negative fraction — the ratio of synergistic versus antagonistic pairs within the community; <u>(b)</u> mean interaction strength — the average correlation coefficient across all possible pairs; and <u>(c)</u> a Cohesion Score, computed as the sum of all pairwise correlation values, where higher values indicate greater predicted community stability. For cross-feeding-based designs, the summary reports the cross-feeding pair fraction (proportion of strain pairs with non-zero metabolic exchange), the mean and total normalized cross-feeding scores, and a qualitative interpretation of overall metabolic complementarity within the community.
5. Data synchronization (*syncom101_correlation_parser.py*): To ensure technical consistency, the correlation parser automatically filters the interaction matrix to match only the strains present in the current functional dataset. It handles the synchronization of strain identifiers, ensuring that the selection engine only considers candidates for which both functional and interaction data are available.

### Evaluation metrics and visualizations

SynCom101 integrates a dual-layered visualization framework that transitions from real-time interactive auditing to the generation of publication-quality static figures.

#### Real-time interactive results (results.py)

Upon generation of SynComs, the platform renders interactive dashboards using the Plotly (a python graphing library, https://plotly.com/python/). These allow users to query specific data points in real-time:

1. Interactive Functional Dot Plots: Maps traits (X-axis) against strains (Y-axis), where color intensity is dynamically scaled to represent trait abundance or gene copy number.
2. Live Coverage Summaries: Comparative bar charts that rank different SynCom variants or optimization strategies based on their total functional scores, allowing for immediate selection of the most efficient design.

#### Figures (Graph_creation.py & export.py)

For formal reporting and publication, the platform includes a dedicated graph-generation engine based on Matplotlib. These figures are optimized for high-resolution export (e.g., PDF/SVG) along with the data for generating these figures:

1. Overlap vs. Improvement Curves: A critical validation plot that illustrates the trade-off between increasing community size and the marginal gain in functional coverage. This plot also visualizes the effect of the “Redundancy Penalty,” contrasting “lean” designs with “robust” ones.
2. Trait Redundancy Histograms: High-resolution distributions showing the frequency of each trait within the community, used to identify functional “bottlenecks” or high-redundancy metabolic pathways.

#### Comparison of SynCom libraries

A unique feature of the results module is the ability to upload user-defined SynCom compositions (CSV format) for comparison. Technically, the platform maps the members of the uploaded community to the master strain-trait matrix to calculate its functional coverage, redundancy, and cohesion metrics using the same parameters applied during the generation phase.

#### Comprehensive Analysis Export Architecture

The Export Module aggregates these different layers. All files can be exported separately, when user select “Complete Analysis Package” the backend triggers the Graph_creation scripts to generate static versions of all session charts. These are bundled into a standardized ZIP archive alongside:

1. *original_data.csv*: original strain-trait data in pickle format
2. *weighted_data.csv*: trait data with applied weights (if available)
3. *algorithm_results.json*: Complete algorithm results
4. *analysis_summary.txt*: Human-readable summary report
5. *selected_strains_trait_matrix.csv*: Filtered matrix with only selected strains and non-zero weight traits
6. *syncom_strain_assignments.csv*: SynCom-strain assignments in 2-column long format (SynCom_ID, Strain_Name). This file is ready to upload to the *Trait Analysis* page for SynCom-trait matrix generation and compatible with other visualization and analysis tools
7. *visualizations folder*: PDF files of all generated plots and figures including <u>a)</u> Performance comparison plots (scatter, box plots) <u>b)</u> Dot plots for all SynComs (ranked by coverage)
8. *“visualizations/data/” folder*: CSV data files used to generate each visualization: Allows users to recreate or customize plots with their own settings.

### Community size optimization (FindSize)

To provide users overview for choosing a proper SynCom size, SynCom101 provides a optimization module (findsize.py) which automates the exploration of the “Size-Function” landscape. The module executes a sequence of SynCom generations across a user-defined range of community sizes (e.g., from 1 to 50 strains) and a step value (e.g., step = 5, tested size will be 5, 10, etc). For each size *k*, the platform runs the parsimony engine with weighted data and records the resulting Functional Coverage Score (*C_k_*), which represents the weighted ratio of traits covered by generated SynCom versus traits involved in microbial candidate pool:

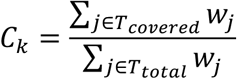

This allows the system to generate a saturation curve that identifies the “elbow point” - the community size where adding further strains yields little improvements in functional coverage.

### Post-selection trait analysis

To bridge the gap between algorithmic output and biological verification, SynCom101 includes a comprehensive *Trait Analysis* module (trait_analysis.py). Unlike the automatic results dashboard, this module operates as a user-driven query engine for the rigorous auditing of specific microbial combinations. The module enables the dynamic construction of functional matrices based on user-defined subsets of data. Users provide a list of specific strain identifiers or generated SynCom IDs for comparison and a list of target traits, the backend performs an *N × M* lookup operation across the master database to generate a tailored Strain-Trait or SynCom-Trait Matrix, allowing for the direct comparison of different community variants.

### Data preparation for benchmarking and case studies

#### Public bacterial collections

The genomes annotated by eggnog-mapper [26,37] and the annotation files were used for benchmarking parser performance. These datasets were preloaded on the website and are available for users on the data input page to merge from different functional annotation types.

1. ProGenomes3 (host plant-associated representative genomes set) [38]: The genomes were downloaded from (http://progenomes3.embl.de/data/habitats/representatives.host_plant_associated.contigs.fasta.gz) and filtered for only complete genomes for follow-up analysis.
2. At-SPHERE (*Arabidopsis* microbiota culture collection) [39]: This dataset represents a comprehensive resource of cultured bacterial isolates from the *Arabidopsis thaliana* leaf and root microbiota.
3. CRBC (Crop root bacterial genome collection) [32]: based on the recent large-scale effort to characterize the crop root microbiome (including rice, maize, and wheat), we filter out the MAGs and only keep the cultured bacterial genomes (4618 genomes). This dataset is large-scale that suitable for benchmarking the tool can be scaled up.
4. metaTraits (https://metatraits.embl.de/) [40]: based on NCBI genus level traits summary files generated from metaTraits database.

#### Custom experimental dataset

For benchmarking the biosynthetic gene cluster families’ matrices from BiG-SCAPE [28] and MEGAMATRIX from bacLIFE [29] we also employed a dataset from a suppressive soil bacterial collection [35]. Similar to preparation for public bacterial collections, the genomes were annotated by eggnog-mapper [26,37], and secondary metabolites potential were generated by antiSMASH [27] and clustered using BiG-SCAPE [28]. Manual curated metallophore biosynthetic gene cluster families (GCFs) were extracted with R (Supplementary Table S1). The genome-based metabolic models (GSMMs) for each genomes were generated with gapseq [36], and the cross-feeding matrix were generated based on produced metabolites and consumed metabolites prediction of the GSMMs (Supplementary Table S2). The cross-feeding matrix and bacLIFE datasets were used in the example study 3.

## Acknowledgements

The work was supported by the European Union via ERC Starting Grant 948770-DECIPHER to M.H.M., the Dutch Research Council (NWO/OCW), as part of the MiCRop Consortium Programme, harnessing the second genome of plants (grant number 024.004.013), and J.J. was supported by China Scholarship Council (Grant No. 202106990013) for her PhD study.

